# PlasmidHostFinder: Prediction of plasmid hosts using random forest

**DOI:** 10.1101/2021.09.27.462084

**Authors:** Derya Aytan-Aktug, Philip TLC Clausen, Judit Szarvas, Patrick Munk, Saria Otani, Marcus Nguyen, James J Davis, Ole Lund, Frank M Aarestrup

## Abstract

Plasmids play a major role facilitating the spread of antimicrobial resistance between bacteria. Understanding the host range and dissemination trajectories of plasmids is critical for surveillance and prevention of antimicrobial resistance. Identification of plasmid host ranges could be improved using automated pattern detection methods, compared to homology-based methods due to the diversity and genetic plasticity of plasmids. In this study, we developed a method for predicting the host range of plasmids based on the random forest machine learning method. We trained the models with 8,519 plasmids from 359 different bacterial species per taxonomic level, where the models achieved 0.662 and 0.867 Matthews correlation coefficients at the species and order levels, respectively. Our results suggest that despite the diverse nature and genetic plasticity of plasmids, our random forest model can accurately distinguish between plasmid hosts. This tool can be used online through Center for Genomic Epidemiology (https://cge.cbs.dtu.dk/services/PlasmidHostFinder/).

**Importance:** Antimicrobial resistance is a global health threat to humans and animals causing high mortality and morbidity, and effectively ending decades of success in fighting against bacterial infections. Plasmids confer extra genetic capabilities to the host organisms through accessory genes, which can encode antimicrobial resistance and virulence factors. In addition to lateral inheritance, plasmids can be transferred horizontally between bacterial taxa. Therefore, detecting the host range of plasmids is crucial for understanding and predicting the dissemination trajectories of extrachromosomal genes and bacterial evolution, as well as for taking effective counter measures against antimicrobial resistance.

## INTRODUCTION

Plasmids are extra-chromosomal DNA sequences that have crucial roles in bacterial ecology, evolution and the spread of antimicrobial resistance (AMR) (1). They are typically circular, self-replicating, transferable and tend to obtain, lose or re-arrange their genetic content rapidly which make them extremely mosaic, diverse and plastic. Plasmids are generally composed of backbone and accessory genes. The backbone includes replication (*rep*) and mobility (*mob*) genes which are relatively conserved amongst the plasmids of the same family (2). These features have also been used to type and compare plasmids that are isolated from different hosts using the replicon and MOB typing (3-5). The accessory genes generally confer selective advantages to the host such as AMR, virulence and metal resistance, increasing host survival under stress conditions despite the metabolic costs that plasmids cause to the host (6). Plasmids also harbor toxin-antitoxin systems and act as parasitic entities (7). Plasmids are often competent horizontal gene transfer vectors, and are able to move from one bacterium to another via conjugation, transduction or transformation causing persistent genetic exchange between bacterial hosts (1, 8).

Plasmids vary in the number and range of taxa they can transfer to, replicate in and be maintained in. They can be roughly categorized as having narrow or broad host ranges (9). The features that determine the host range capacity of plasmids are not fully understood yet, but origin of replication, replication initiation dependencies, and origin of transfer are known to be important for host range (9).

Plasmid host ranges can be determined empirically by testing potential hosts *in vitro* (10, 11). However, sequence-based approaches can be used for plasmid host range prediction, which is more practical compared to the empirical methods in terms of turn-around times and usage of laboratory resources (11). Previous studies have attempted to predict plasmid host ranges by comparing oligonucleotide composition of plasmids and chromosomes (1, 9, 10, 12-14). Narrow host range plasmids are expected to have similar oligonucleotide compositions with the host organism due to plasmid sequence amelioration *e*.*g*. adaptation to a preferred host codon usage (12). However, this method falls short when predicting broad host range plasmids because of plasmids can often transfer to distantly related hosts (9, 12).

Previously developed plasmid identification tools such as PlasmidFinder and PlasFlow have been developed to determine plasmid hosts (3, 15). PlasmidFinder identifies plasmids in whole genome sequences by searching against plasmid replicon sequences from the *Enterobacteriaceae* and Gram-positive species. This alignment-based tool identifies plasmids from these taxa with high accuracy by indicating a source organism based on the best matching replicon. PlasFlow was developed using deep neural network and trained by the *k*-mer counts of fragments at least 1,000 nucleotides in length, and it can detect plasmid hosts at the phylum level. To our knowledge PlasFlow tool is not currently maintained. Recently, Redondo-Salvo et al. (16) developed an automated plasmid classification tool by assigning plasmid taxonomic units (PTUs) using total average nucleotide identity.

Machine learning, a form of artificial intelligence, has been utilized in recent years to understand various biological systems by detecting the linear and non-linear correlations between input and output data (17). It has been used to predict phenotypes and structures in nature, and it has the potential to discover unknown features such as novel AMR genes (18-20). In this study, in order to better predict plasmid hosts and infer plasmid host ranges, we developed a set of random forest-based machine learning models for predicting plasmid hosts at several bacterial taxonomic levels.

## MATERIAL AND METHODS

### Data set

We downloaded all of the available (10,863) plasmids and corresponding metadata from the Pathosystems Resource Integration Center (PATRIC) (21) in September of 2020. Metadata for each plasmid included the origin of the plasmid and other relevant information such as different database accession numbers, collection date and place, genomic length and features. Four plasmids did not have host information and were removed. The remaining plasmid host information was reported from genus to strain level by the PATRIC database. In total, 1,662 different genus and species level hosts were detected in the plasmid metadata. When fewer than five plasmids had a given host at a given taxonomic level, they were removed. In total, 1,296 under-represented hosts and corresponding plasmids were removed from the dataset to improve the robustness of the models. From the remaining 366 hosts, seven were removed for lacking species annotation. Therefore, we generated machine learning models with 8,519 plasmids and 359 corresponding hosts with species level taxonomy information. The species-level plasmid hosts were assigned to the higher taxonomy levels such as genus, family and order using the NCBI Taxonomy information by the Python ete2 package (version:2.3.10) (22, 23).

### Distance tree

The diversity of the plasmids was measured using an oligonucleotide *k*-mer-based distance tree. The plasmid sequences were indexed using the KMA tool (version: 1.3.9) (24) with the following parameters: -NI -Sparse TG. Next, the *16*-mer Hamming distances were calculated. The distance tree was generated using the CCPhylo tool (version: 0.2.2) using the Neighbor-Joining method (25). The distance tree was visualized using iTOL (version: 4) (26).

### *K*-mer counts

The plasmid genomes were sub-sampled using overlapping *k*-length nucleotides (*k*-mers) and counting the occurrence of every sub-sequence. *K*-mer counting is a well-studied method for analyzing sequence data (27). The sub-sequence size *k* is a critical parameter as the sub-sequences yield various information depending on the size. While short *k*-mers provide information regarding the sequence content, long *k*-mers are informative in detection of unique sequence patterns. We analyzed plasmid genomes using three different *k*-mer sizes: 5, 8 and 10 nucleotides. Counts were calculated using KMC (version: 3.0.0) (28) with the following parameters -fm, -ci1, -cs1677215. These parameters inform the tool regarding the input data format and the minimum and maximum thresholds for the *k*-mer occurrences.

### Detection of duplicates

To eliminate possible duplicates from the plasmid collections, we compared the *8*-mer counts of the plasmids to each other. Plasmid pairs with identical *8*-mer counts were treated as duplicates and merged in the dataset. When the pairs had differing host information, the additional hosts were incorporated in the metadata.

### Sequence length, GC content and codon usage calculation

To capture the plasmid genome characteristics, we calculated the total length of the sequence, GC content and codon usage. Sequence length was calculated by taking all nucleotides into consideration, including ambiguous bases. GC content was calculated by taking the ratio of the total number of cytosine and guanine nucleotides to all nucleotides. Codon usage was determined as the relative frequencies of codons in a coding region which was detected using Prodigal (version: 2.6.3) with default parameters (29).

### Model generation and cross validation

For each *k*-mer size, a matrix was generated from *k*-mer counts where the rows represent each plasmid, the columns represent each *k*-mer and the entries represent the *k*-mer counts. Additionally, a merged matrix was generated by combining the *8*-mer count matrix with the genome length, GC content and codon usage information.

In this study, we generated multi-label models that are able to predict multiple hosts per plasmid. Each label corresponds to a plasmid host and encodes a binary value, with “1” corresponding to being a host. These plasmid hosts were predicted at different taxonomy levels such as species, genus, family and order, where we built separate models per taxonomic level. We used random forest to build the classifiers, which provides robust and interpretable predictions based on decision trees and has been explored in many other classification studies (18, 30-32).

Model parameter tuning and validation was performed using the plasmid data, where the data were split into training, testing and hold-out datasets. The training and testing datasets were used for parameter tuning, and the hold-out dataset used for monitoring possible overfitting. Random forest was implemented using *ensemble*.*RandomForestClassifier* from the Python Scikit-learn package (version: 0.20.4) (33). The model parameters were tuned in the five-fold cross validation loop using the random grid search method from Scikit-learn that iterated 100 times; n_estimators, max_features, max_depth, min_samples_split, min_samples_leaf and bootstrap were the parameters tuned (Table S1). These parameters were responsible for the number of trees in the forest, the number of features required for the split, the maximum depth of the tree, the minimum number of samples for splitting, the minimum number of samples required for the leaf, and bootstrapping of samples, respectively. Tuning was conducted using an *8*-mer matrix at the genus level and then applied to the other taxonomic levels and *k*-mer sizes. The detected optimal parameters were n_estimators = 1,000, max_features = “auto” which is square root of the number of features, max_depth = 50, min_samples_split = 2, min_samples_leaf = 1, and bootstrap = False. The class_weight parameter was set to “balanced” to weight the inputs based on the class frequencies to prevent the biased predictions due to the imbalanced classes. The random forest model was utilized with the *multiclass*.*OneVsRestClassifier* from the Python Scikit-learn (version: 0.20.4) package (33) which fits one label at a time and improves the interpretability of the models.

Using the tuned parameters, the random forest model was trained and tested five times using the *k*-fold cross-validation method, where different datasets were tested each time. The ensembled cross-validation model was applied on the hold-out dataset which was not part of the training or testing. Model performances were measured using the area under curve (AUC), macro F1 score, Matthews correlation coefficient (MCC), and the confusion matrix using Scikit-learn (33). Sensitivity and specificity were also calculated from confusion metrices to measure the ability of model to identify hosts. The class probability threshold of 0.5 was applied to calculate the performances of macro F1, MCC, and confusion matrix. Since it is a multi-label problem, all of the predictions for all possible labels were collected into one data container, and one prediction performance was calculated per model instead of the number of labels. The test and hold-out dataset performances were reported. The test performances were calculated by averaging five model performances from the cross-validation, and reported with standard deviations. The hold-out performances were the ensemble model performances from averaging the cross-validation model predictions.

To test whether the plasmid host model predictions were significantly different, a t-test was performed using *stats*.*ttest_*ind from SciPy (version: 1.2.2) (34). Moreover, possible correlations were detected using Spearman’s correlation coefficient using *stats*.*spearmanr* from SciPy (version: 1.2.2).

### Clustering plasmids

Since similar plasmids are likely to be hosted by the same organisms, we clustered the plasmids based on *k*-mer sequence similarity using KMA index (version: 1.3.9) with the following parameters: -k16, -Sparse and -NI (24). KMA clusters the sequence for a given similarity threshold using *16*-mers and the Hobohm-1 algorithm (35). We clustered the plasmids using three different *k*-mer query and template similarity thresholds at 90%, 80% and 50%. By dividing the clusters into training, testing and hold-out sets; similar plasmids were kept in the same partitions. This forced the models to learn sequence characteristics spanning larger genetic distances and was intended to help improve the generalizability of the models.

### Random fragments

Partial sequences might be informative for predicting hosts and better reflect actual data in incomplete plasmid assemblies. Therefore, random fragments of 500, 1,000, and 1,500 nucleotides were sub-sampled from each plasmid sequence to build prediction models from the partial sequences. The sub-sampling process was repeated randomly ten times for each plasmid. Plasmids shorter than the given fragment size were excluded from the study and separate models were built per fragment size. Matrix files and models were constructed as described above, using *k*-mers that were 5 and 8 nucleotides in length. *10*-mers were not utilized due to the heavy computational requirements.

### Validation of the plasmid host prediction models

The plasmid host models that were trained with the PATRIC plasmids at four different taxonomic levels were validated using plasmids from the National Center for Biotechnology Information (NCBI) Reference Sequence database (RefSeq) (36). A total set of 30,349 NCBI plasmids were downloaded from the NCBI RefSeq database and filtered. NCBI offers a larger plasmid collection than PATRIC, yet some of the plasmids are identical. Therefore, only the plasmids that had not yet been integrated into PATRIC as of January 2021, *i*.*e*., plasmids that are only present in the NCBI RefSeq database were included. Further, we eliminated duplicates from the NCBI validation dataset by comparing *k*-mer counts, and filtered based on the source organism, completeness and NCBI’s automatic taxonomy check. Moreover, plasmids with labels that are not included in the PATRIC training data were further removed from the NCBI validation data. The remaining plasmids with species level host information recognized by NCBI Taxonomy were tested against the plasmid host models that were trained on the PATRIC collection. The validation performances were reported in AUC, macro F1, MCC and the confusion matrix. The class probability threshold of 0.5 was applied to calculate the performances of macro F1, MCC and confusion matrix.

### Comparison to PlasmidFinder

We compared our species model performance to PlasmidFinder for the *Enterobacteriaceae* species that are present in the PATRIC hold-out dataset (3, 37). Moreover, the hold-out plasmids that present in the PlasmidFinder database were excluded from this comparison. The PlasmidFinder tool (version: 2.1.1) was performed with the default parameters using the PlasmidFinder database downloaded in July 2021.

## RESULTS

### Plasmid host prediction performances for the PATRIC hold-out dataset

In order to develop models for predicting the host organisms of plasmids, a total of 8,519 plasmids with at least species level host information were downloaded and curated from the PATRIC database and included in this study (Table S2). These plasmids originate from 359 species belonging to 174 genera, 93 families, and 50 orders (Figure S1, Table S3). Most of the plasmids in the collection come from the orders Enterobacterales, Bacillales, and Lactobacillales, which comprise 55.6% of the hosts in the data set (Figure 1).

**Figure-1:**
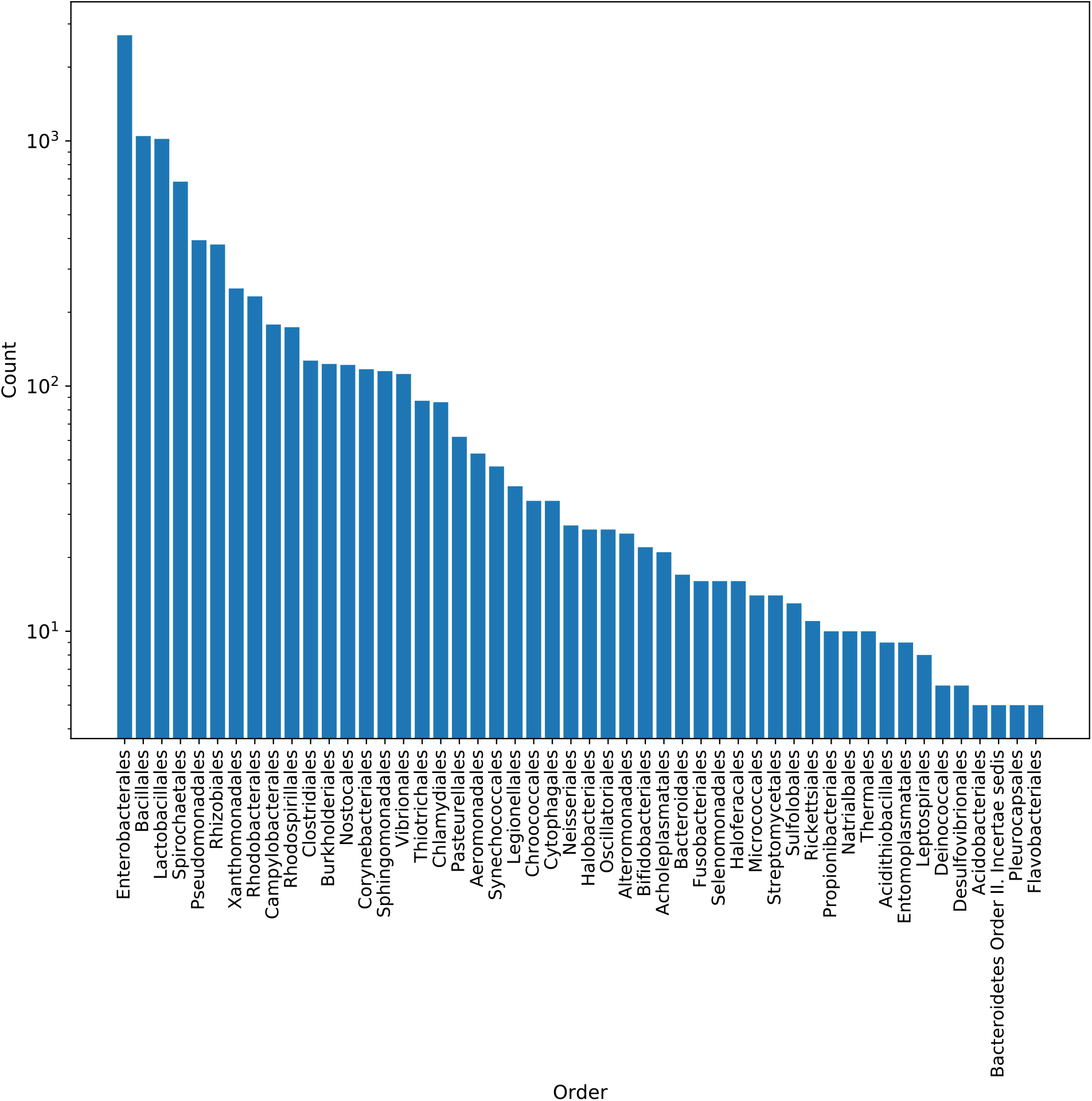
The plasmid host distribution at the order level in the PATRIC dataset. The PATRIC plasmid collection was dominated by the Enterobacterales, Bacillales and Lactobacillales orders which make up 55.6% of the plasmid hosts.

To predict the taxonomic label of the host organism, machine learning models were trained using nucleotide *k*-mer counts from the plasmids. The predictions were carried out using *5, 8* and *10*-mers, since the short and long *k*-mers might provide different types of information to the models. For example, *5*-mers do not usually appear in the plasmid genome uniquely, and instead provide the models with information regarding the profile of oligonucleotide frequencies for each plasmid. On the other hand, the longer *k*-mers, such as *8*- and *10*-mers, usually occur uniquely in a given plasmid, and offer counts of unique sub-sequences. Moreover, *k*-mer distributions are subject to changes based on the sequence size.

Using each *k*-mer size, random forest-based classifiers were built to predict host taxonomy from order to species levels. The model based on the *5*-mer counts has 0.655 MCC for predicting the plasmid host species, and this was moderately higher, 0.662 and 0.680 MCC, for *8*-mers and *10*-mers, respectively (Figure 2, Table S4-S6). At the order level, the model performances achieved 0.899 MCC for *5*-mers, 0.867 MCC for *10*-mers and *8*-mers (Figure 2, Table S4-S6).

**Figure-2:**
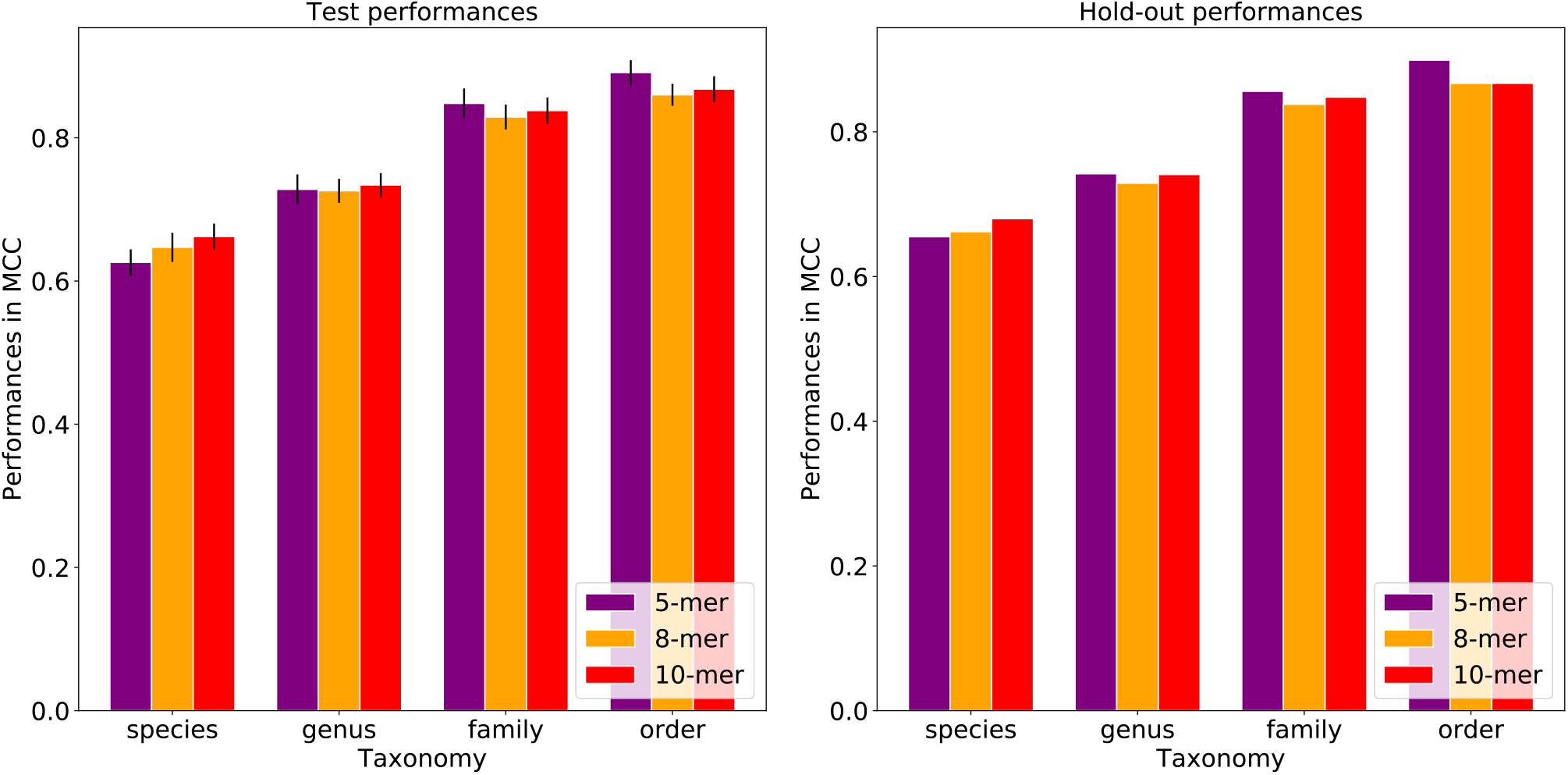
Host prediction performances by *k*-mer size for the test and hold-out data sets. Each bar represents the model performance per taxonomic level. While the test performances were reported with standard deviations, the hold-out performances do not have standard deviations as the five models were combined and a single performance was calculated. The plots show that the prediction performances vary when using different *k*-mer sizes. *5*-mers yield the highest MCC at higher taxonomies, while *10*-mers yield the highest MCC at lower taxonomies. The model performances generally increased from the species to order level for all the *k*-mer sizes.

By increasing the *k*-mer size from five to ten, the prediction performances increased 3.8% in MCC at the species level but decreased 3.6% at the order level. Although the fluctuations in the performances are not significant according to the paired t-test (p-values [0.404, 0.883] > significance threshold 0.05). To limit computational needs, we used the *8*-mers, but not *10*-mers, to build input matrices for all sub-sequent analyses. Overall, the plasmid host prediction models have low sensitivity (true positive rate) and high specificity (true negative rate) where the lowest sensitivity was detected at the species level compared to other taxonomy levels where sensitivity falls into the range between 0.493 and 0.761.

The number of the false negative predictions increased inversely with the presence of the hosts in the input data (Figure 3). Moreover, this correlation was significant at the species level (Spearman’s correlation coefficient of 0.545, p-value 0.00 < significance threshold 0.05). These findings suggest that the host classification becomes more challenging at the species level, and the model performances improve proportionally to the host representations in the training data. In addition to the false negatives, false positive predictions made by the model (Figure S3-S4) were frequently phylogenetically close to the actual hosts. For instance, the model frequently predicted *Escherichia coli* hosts instead of the *Salmonella enterica* and *Klebsiella pneumoniae*, where all of them belong to the *Enterobacteriaceae*.

**Figure-3:**
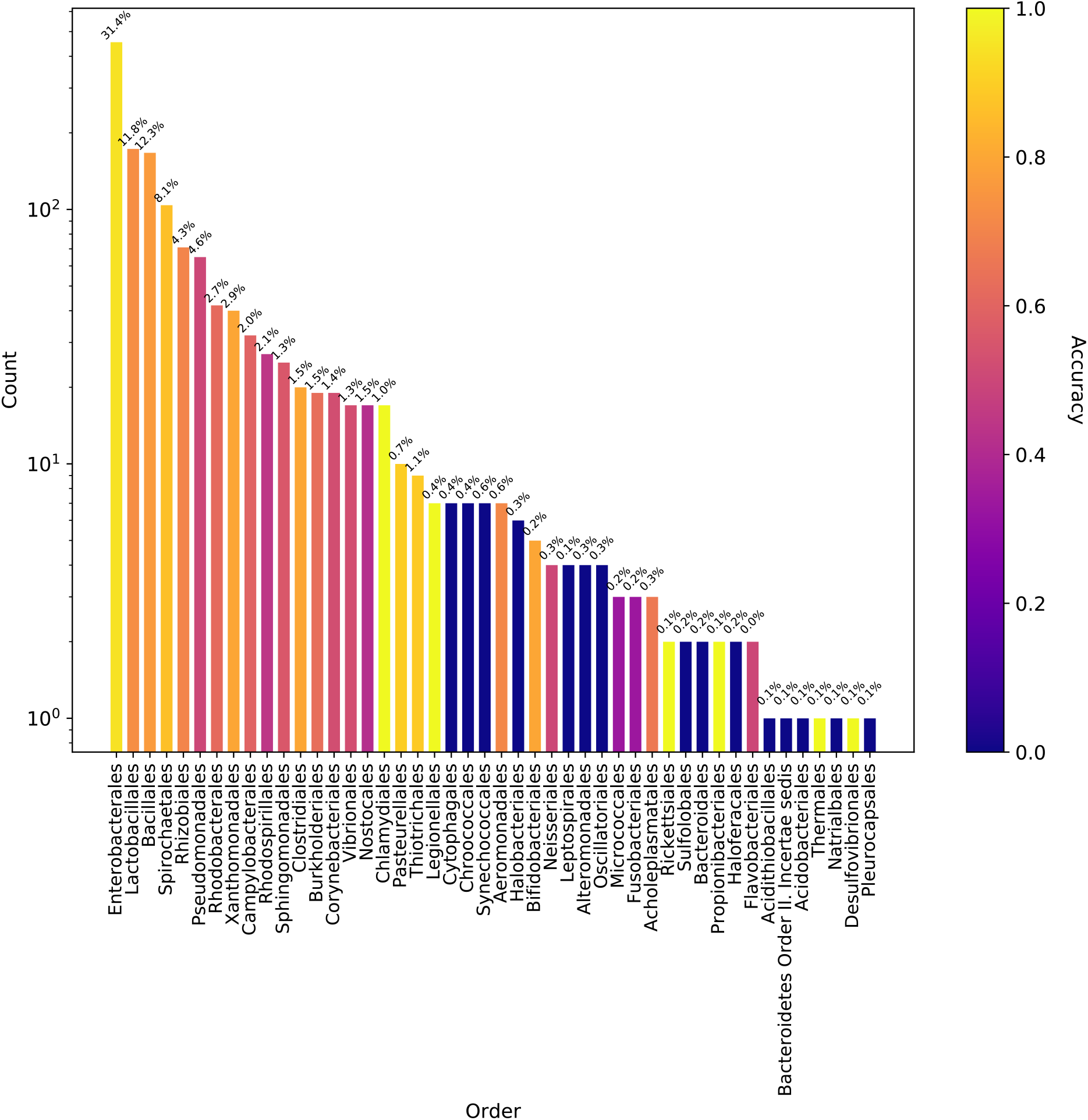
Model accuracy for the PATRIC plasmids tested with the whole model. Each bar shows the number of bacterial orders in the hold-out data and corresponding model accuracy was color coded. The percentage on the top of each bars shows the percentage of bacterial orders in the training data.

In an attempt to improve the *8*-mer model performances, we combined the *k*-mer frequencies with the information in nucleotide compositions of plasmid sequences including, plasmid size, GC content and codon usage. These additional features yielded an approximately 0.6%-1.6% increase in the MCCs of the models (Figure S2, Table S7); however, this improvement was not significant according to the paired t-test (p-value 0.892 > significance threshold 0.05). Therefore, the following analyses were carried out without these additional features.

To understand the impact of plasmid sequence similarity on the model performances, the plasmid genomes were clustered based on the *k*-mer similarity using KMA. The plasmids belonging to the same cluster at a given *k-*mer similarity threshold were kept in the same training, testing or hold-out dataset. When the *k-*mer similarity decreased to 80%, thus making the clusters more diverse, the model performances decreased in MCC between 7.7% to 29.8% depending on the taxonomic level (Figure 4, Table S8). The performance decrease shows that the similarity between the training and testing data has an effect on the host predictions, especially at the lower taxonomic levels, although, the model can still be generalized to distant sequences.

**Figure-4:**
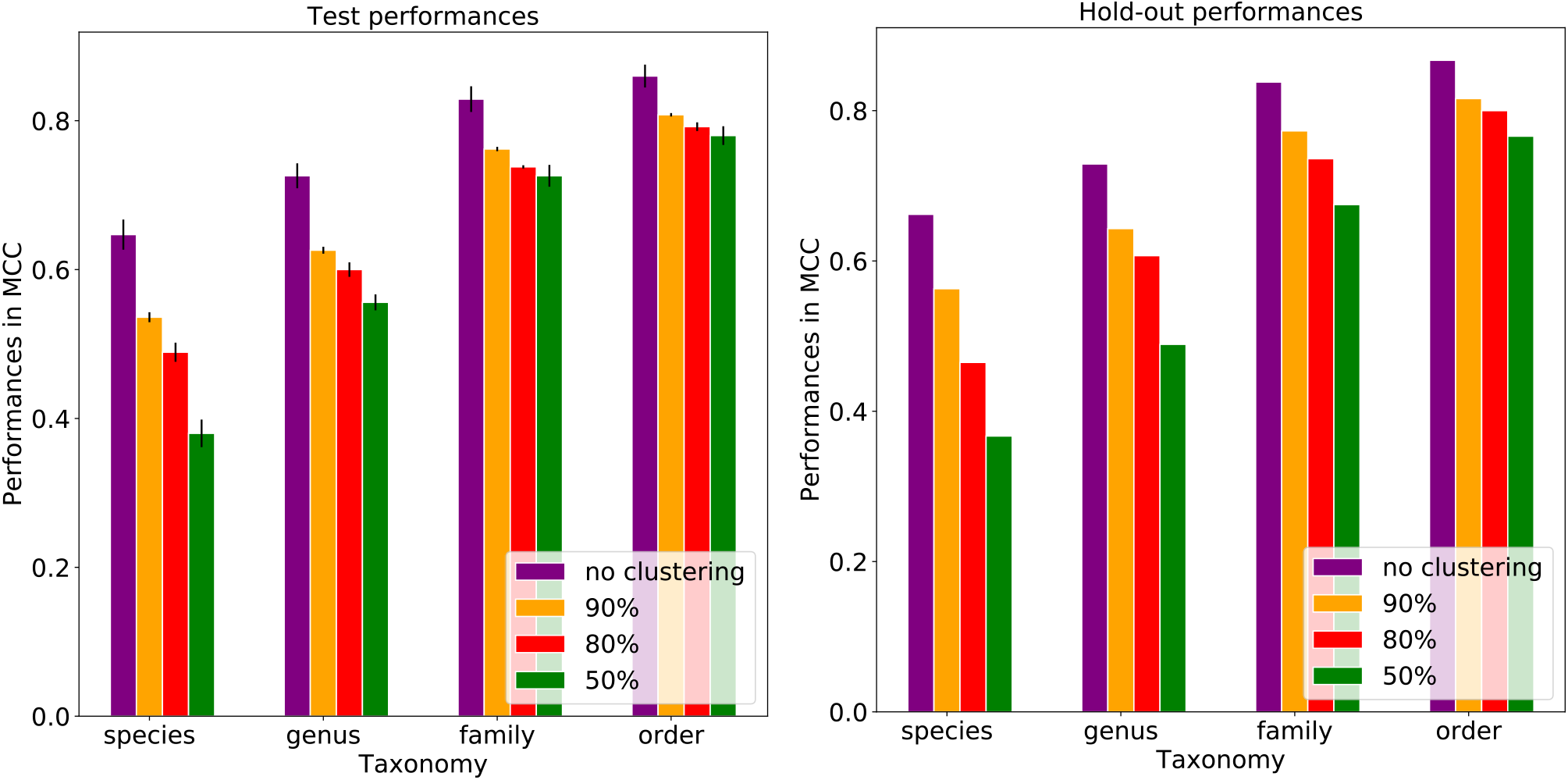
The effects of clustering plasmids at different *k-*mer similarity thresholds on the plasmid host predictions using *8*-mers and different taxonomic levels. Each bar represents the model performance per taxonomic level and each error bar represents standard deviations across folds. The plot shows the influence of plasmid sequence similarity on prediction performances in MCC from the species to order level. The plots suggested that the prediction models pick up sequence similarity mostly at lower taxonomic levels. When the dissimilarity was increased between the training, test and hold-out datasets by applying the 80% *k*-mer similarity threshold; 7.7%-29.8% in MCC performance loss were observed for the hold-out data.

### Plasmid host predictions with random fragments

Due to the fragmented nature of plasmid assemblies that results from the difficulty in assembling plasmids from the short reads, we wanted to develop random forest models that can make predictions from incomplete sequences. To do that, we trained and tested our plasmid host prediction models with random fragments of plasmid sequences. Fragments of 500, 1,000 or 1,500 nucleotides were randomly sampled from each assembled plasmid sequence over ten rounds. By sampling multiple times, we attempted to introduce various regions of the plasmid sequences to the models. The fragment model that was trained with the 500 nucleotide fragments using *5*-mers reached 0.426 MCC for the species model and 0.674 for the order model (Figure 5, Table S9). When the same fragments were sub-sampled into *8*-mers, the species level model had MCCs of 0.489 and 0.686 MCC for the species and order levels, respectively (Figure 5, Table S10). By increasing the fragment size from 500 to 1,000 nucleotides, the model performances increased 8.2%-10.7% in MCC with the *5*-mers and 6.3%-10.1% in MCC with the *8*-mers (Figure 5, Table S9-S10). When the fragment size increased from 1,000 nucleotides to 1,500 nucleotides, the model performances increased 4.8%-5.1% in MCC with the *5*-mers and 3.3%-1.7% in MCC with the *8*-mers (Figure 5, Table S9-S10). The fragment models reached their highest performances using 1,500 nucleotide fragments and *8*-mers as the features, where the MCCs were 0.537 and 0.768 for the order and species model, respectively (Figure 5, Table S10).

**Figure-5:**
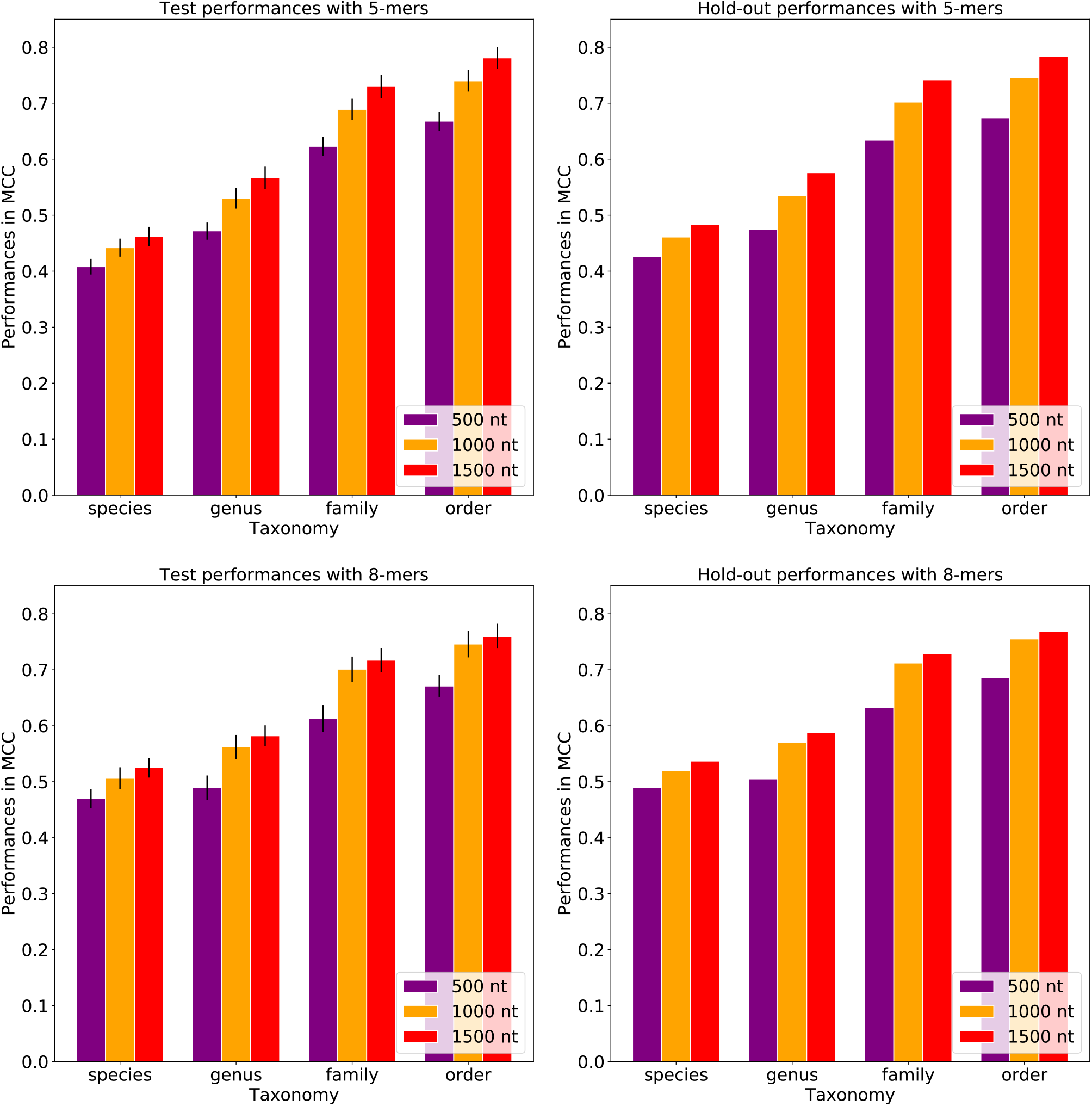
The fragment model performances for the *5*-mer and *8*-mer models. The fragment models were trained with either 500, 1,000, or 1,500 nucleotide (nt) fragments that were sub-sampled from the PATRIC plasmids. The bar plots show the test and hold-out performances for the *5*-mers and *8*-mers in MCC. The error bars represent standard deviations. The best performing model was trained with the 1,500 nucleotide fragments using *8*-mers.

### Validation of the plasmid host prediction model with the NCBI validation dataset

To validate the plasmid host prediction models, we used plasmids in the NCBI RefSeq collection that are not present in our training, test or hold-out datasets. Overall, 7,670 bacterial plasmid sequences with taxonomic metadata were included in this analysis (Table S11-S12). As in the PATRIC database, the NCBI validation data is also dominated by the few major orders such as Enterobacterales, Lactobacillales and Pseudomonadales, which make up approximately 76% of the data set (Figure 6).

**Figure-6:**
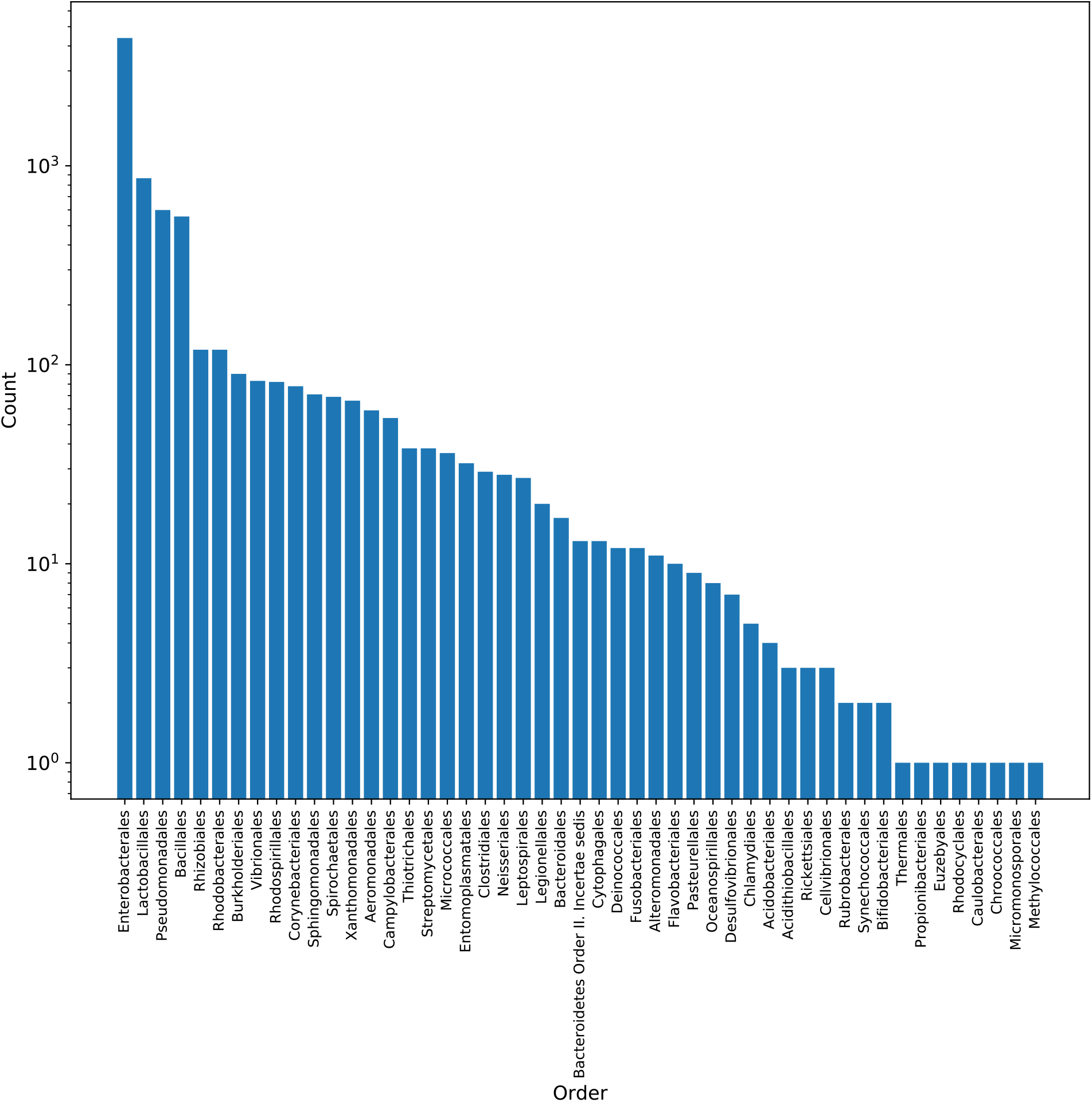
The plasmid host distribution in NCBI validation dataset. The validation dataset was dominated by the Enterobacterales, Lactobacillales and Pseudomonadales, which make up 76% of the NCBI plasmid hosts.

When the whole model (trained with *8*-mers of the PATRIC training set) was tested with the NCBI validation data, the ratio of the correct and wrong predictions was shown in Figure 7. Our plasmid host prediction model has relatively low sensitivity (0.483) and a high specificity (1.0) at the species level (Table S13), similar to the results shown above. Moreover, when the NCBI validation data were tested with the random model generated by shuffled labels, the model performance dropped to 0.028 MCC at the species level. This suggests even though the sensitivity is low, the model has adequate generalizability which is far from being random.

**Figure-7:**
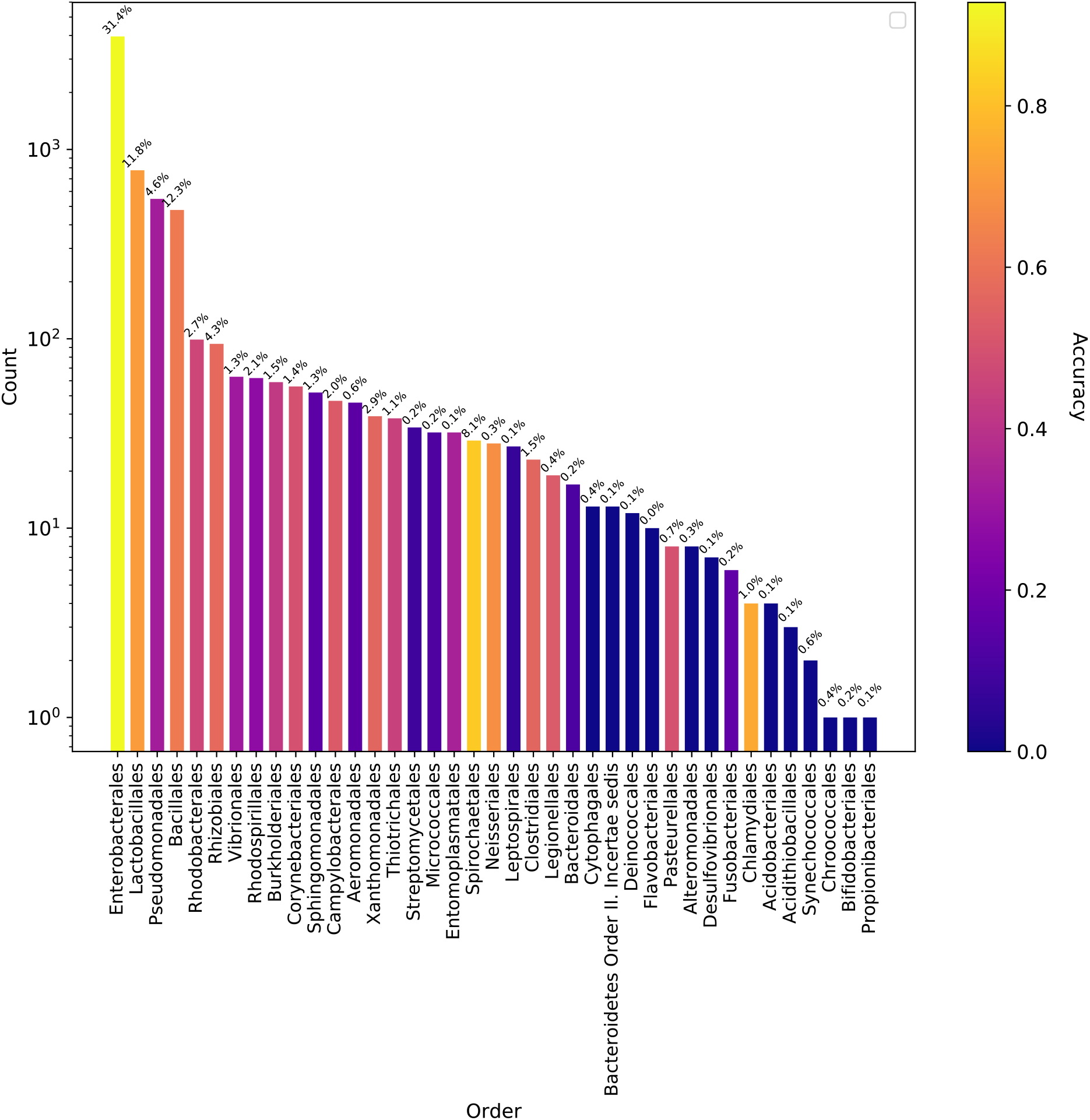
Model accuracy for the NCBI plasmids tested with the whole model that was trained with the PATRIC dataset. Each bar shows the number of bacterial orders in the validation data and corresponding model accuracy was color coded. The plot showed that the accuracy of the models changed roughly according to the availability of the host organisms in the training data which was indicated on top of the bars.

Because the NCBI collection contained many short plasmid sequences, we filtered it based on the sequence size. Overall, plasmid sequences equal or greater than 5,000 bp performed 43% better than plasmid sequences less than 5,000 bp in terms of MCC at the species level. However, this performance gap reduced to 1% at the order level (data not shown). This means that the plasmid host range model accuracy improves with longer plasmid sequence length at lower taxonomic levels.

Similarly to the predictions for the hold-out dataset, the plasmid host prediction model predicted additional hosts for 499 plasmids in the NCBI validation dataset (Figure S5). Furthermore, the model frequently erroneously predicted *Escherichia coli* as being the host instead of its close relatives, *Salmonella enterica* and *Klebsiella pneumonia*, both in the hold-out and the validation results.

The fragment-based models were also validated using the NCBI dataset. The 7,670 NCBI plasmids were randomly sub-sampled into 500, 1,000, and 1,500 nucleotide fragments, and each plasmid was randomly sampled ten times per fragment size. Similar to the PATRIC results, the fragment models reached the best performances (0.485 MCC at the species level and 0.778 MCC at the order level) for the NCBI validation data with the 1,500 nucleotides fragment size and *8*-mers (Figure 8, Table S14-S15).

**Figure-8:**
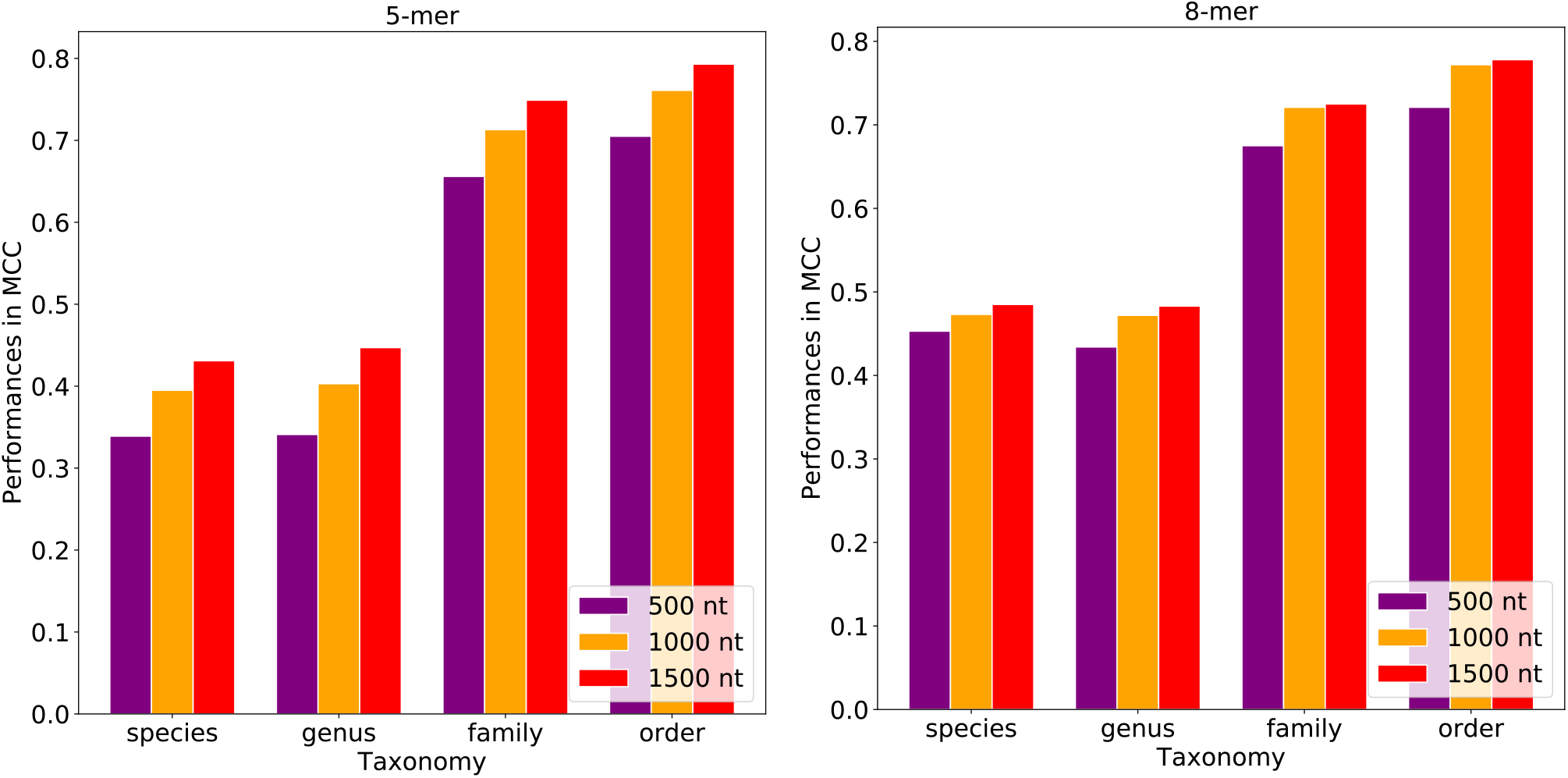
The fragment models were validated with the NCBI plasmids. The fragments models that trained with the 500, 1,000 and 1,500 nucleotide (nt) fragments from the PATRIC plasmids were validated with the fragments that sub-sampled from the NCBI plasmids. Similar to the hold-out results, the best performance was obtained with the 1,500 nucleotide fragments and *8*-mers.

### Comparison to PlasmidFinder

The PlasmidFinder tool uses an alignment-based strategy to identify plasmid sequences, and can often provide host information when it is available. We used 391 *Enterobacteriaceae* plasmids in the PATRIC validation data that were not already part of the PlasmidFinder database to compare the output of PlasmidFinder and our machine learning models. Overall, PlasmidFinder correctly identified 304 of the sequences and did not predict anything for 87 sequences. Our whole-plasmid-based *8*-mer model successfully classified 229 of the plasmid hosts, nothing predicted 121 and falsely predicted 41 of them at the species level. We then randomly sampled the 391 plasmids into 1,500 nucleotide fragments and compared PlasmidFinder with our *8*-mer based model based on 1,500 nucleotide fragments. Overall, PlasmidFinder is able to identify 228 out of 3,910 fragmented plasmid sequences. However, none of the returned matches contained plasmid host information. Our fragment-based model predicted a host for 1,927 of the fragmented plasmid sequences correctly, nothing predicted 1,309 and falsely predicted 674 of them. Compared to our machine learning model, the alignment-based PlasmidFinder tool provides accurate predictions for the *Enterobacteriaceae* species when a sequence match is available and plasmids sequences are mostly complete. When the plasmids are fragmented, the machine learning strategy becomes more advantageous.

### The web-server

The plasmid host prediction models that were trained using *8*-mers from whole plasmid sequences can be used online on the Center for Genomic Epidemiology (https://cge.cbs.dtu.dk/services/PlasmidHostFinder/). This web-server tool accepts one FASTA file at a time and provides an output file containing the predicted plasmid host range at the selected taxonomic level from species to order. The web-server tool enables two model options, fast and slow, with various class thresholds. The slow model uses all five cross-validation models to make a final decision on the plasmid host range. The fast mode uses only the first cross-validation model out of five to predict the plasmid host range. Therefore, one can expect to obtain more confident predictions with the slow model.

## DISCUSSION

In this study, we built random forest models that can predict plasmid hosts and host-ranges at taxonomic levels between species and order; these models achieved accuracies from 0.662 to 0.867 MCC. The model performs better at higher taxonomic levels, with ‘order’ level being the best. We observed that the *k*-mer size does not have a significant influence on the prediction performances. Among the three *k*-mer sizes, we chose to build our prediction models with *8*-mers since it provides robust predictions at all taxonomic levels with less computational effort than *10*-mers. Moreover, we tried to improve the host range predictions with the additional genome features such as plasmid size, GC content and codon usage but the increase in the prediction performances was negligible. We validated our models using an independent dataset from the NCBI RefSeq. These performances were comparable with our previous test and validation results. In addition, to assess the utility of this approach with partially assembled plasmid sequences, we generated models for 500, 1,000, and 1,500 nucleotide fragments, and even the smallest fragments of 500 nucleotides have sufficient information for the identification of plasmid hosts.

### Machine learning

We observe that the robustness of the models is dependent on the quantity, quality and accuracy of the input and output data. In this study, the plasmid host prediction models might suffer from incomplete metadata despite our best efforts. The plasmid data and corresponding plasmid hosts were retrieved from the PATRIC database. However, the PATRIC dataset is likely to contain some plasmids with incomplete host range information. This issue might have an effect on robustness of the models, but is most likely to have a minor impact due to the relatively large input dataset. Nevertheless, some of the false positive predictions might be the consequence of incomplete metadata. Some of the other false positives might potentially be new discoveries relating to plasmid transmission in diverse hosts, although this theory should be validated experimentally.

Plasmid genomes are extremely plastic (38). Accessory genes vary in their presence or absent from the plasmids, which makes plasmid host prediction a complicated task. In order to understand the impact of the genome similarity on the plasmid host model learning, we clustered the plasmids for a given similarity threshold. By keeping the similar plasmids in the same training, test or hold-out datasets, the learning from the sequence similarity was minimized since the similar plasmids tend to have the same hosts. This clustering approach caused less accurate results than the baseline model. These results suggest that sequence similarity has an impact on the model learning. Therefore, to boost the model performances, the training data should be updated regularly to increase the input diversity when more plasmid data is available. In addition to the sequence similarity, host related signals from the relatively conserved regions of the plasmid sequences such as *rep* or *mob* genes are likely learned from the model. Further analysis of the top model features may help to validate or elucidate genetic features involved in transmission, especially in less studied taxonomic groups.

The model performances were evaluated using several performance measurements including AUC, MCC, and macro F1. The AUC and MCC performances were not always correlated and caused different conclusions in some cases such as in the random forest model with the clustered plasmids. The reason for this discrepancy may be the applied class thresholds. AUC uses a range of thresholds to measure the model performances and does not require a defined class threshold. In contrast to AUC, the MCC and macro F1 calculation require predictions instead of probabilities. Therefore, a defined class probability threshold is needed for converting probabilities to predictions. This threshold was set to 0.5 for all the models. But, this threshold might not be the ideal threshold for some of the models, particularly for the imbalanced classes (39). For instance, the species level prediction model has a lower sensitivity (0.493) compared to its specificity (1.0). In other words, the model failed to predict some of the hosts (Figure 3,7), and the majority of the failed predictions were the result of having no positive class predicted for the tested plasmid due to no predictions being above or equal to the class probability threshold of 0.5. Therefore, adjusting the class probability threshold could be a way to improve the model.

### False positives or unknown hosts

The machine learning models have the potential for discovery of unknown correlations between the input features and predicted phenotypes. For example, in previous studies, novel AMR genes were reported using the machine learning models (18, 19). In our case, machine learning might be useful for discovering unknown plasmid hosts. We explored the false positives as in: 1) the model was not able to predict the actual hosts, but predicted false positives (Figure S3), 2) the model predicted multiple hosts including the actual hosts and false positives (Figure S4). These cases should be investigated further as these could happen due to two reasons: the model might pick up noise, or the falsely predicted host might actually be a host in nature. Thus, a portion of the false positives might be the actual hosts which are not discovered before, but machine learning gives the opportunity for it *in silico*. To prove that they are potential hosts would require *in vitro* experiments to test the stability of the plasmid in these bacteria.

### Fragments

The fragment-based model performances vary based on the fragment and *k*-mer sizes. We obtained the best performances for the hold-out dataset with the 1,500 nucleotide fragments using *8*-mers. The fragment size and model performances changed proportionally because the longer fragments are providing more information to the models. This correlation between the model performances and fragment size might be the consequence of the mosaic nature of plasmids. Genes located on plasmids could originate from different organisms and random sampling of these acquired genes might cause false predictions. Moreover, as the plasmids were not aligned prior to the fragmentation, the genetic content of fragments that sub-sampled from different plasmids did not match. Therefore, we expect the model learning the fragment structures instead of the unique patterns.

## Conclusion

We built random forest models and incorporated them in PlasmidHostFinder tool to detect plasmid hosts and host-ranges at various taxonomic levels from species to order with the performance of 0.662 MCC to 0.867 MCC. PlasmidHostFinder can detect a diverse range of hosts for 359 species, 174 genera, 93 families and 50 orders with high accuracy in spite of the mosaic, diverse nature and genetic plasticity of plasmids. The approach described in this study helps to fill a gap in our ability to predict plasmid hosts, particularly in understudied taxa, or when plasmid sequences are fragmented.

## DATA AVAILABILITY

The Python 2.7.15 scripts that used in this study are available on Bitbucket (https://bitbucket.org/deaytan/plasmid-host-prediction/src/master/). The web-server is available on Center for Genomic Epidemiology (https://cge.cbs.dtu.dk/services/PlasmidHostFinder-1.0/). All the PATRIC and the NCBI RefSeq sequences and corresponding metadata can be accessed through the PATRIC (https://www.patricbrc.org) and NCBI (ftp://ftp.ncbi.nlm.nih.gov/refseq/release/plasmid/) resources, respectively.

## ACKNOWLEDGMENTS

We would like to thank Frederik Teudt for his patiently and carefully testing the PlasmidHostFinder tool.

This work was funded by the Novo Nordisk Foundation (grant NNF16OC0021856: Global Surveillance of Antimicrobial Resistance awarded to FMA and OL).

The funding bodies did not play any role in the design of the study or writing of the manuscript, nor did they have any influence on the data collection, analysis or interpretation of the data, or the results.

The authors declare no conflicts of interest.

